# A portable cortical evoked potential operant conditioning system (C-EPOCS): System development

**DOI:** 10.64898/2026.01.08.698448

**Authors:** Disha Gupta, Jodi Brangaccio, Helia Mojtabavi, N Jeremy Hill

## Abstract

This study presents customizations and evaluations aimed at adapting the Cortical-Evoked Potential Operant Conditioning System (C-EPOCS) into a portable, user-friendly platform for real-time neurofeedback applications. A primary goal was to simplify the component-heavy setup by integrating electroencephalography (EEG) and electromyography (EMG) data acquisition into a single system—while still supporting cortical and muscle response assessment and real-time feedback.

One key limitation of portable biosignal acquisition systems is their typically lower sampling rates (e.g., 300–600 Hz) compared to high-resolution systems (e.g., 3200 Hz), which are commonly used for detecting transient responses such as the H-reflex and M-wave. In a C-EPOCS setup, these responses are useful for determining the target stimulation intensity and minimizing inter-session variability in effective afferent excitation.

We evaluated whether lower-resolution EMG signals could still support the generation of H-reflex and M-wave recruitment curves for determining target stimulation intensity. Results showed that while EMG sampled at ∼600 Hz and ∼300 Hz produced greater dispersion in recruitment curve data—particularly at 300 Hz—they still yielded comparable estimates for stimulation intensities that elicit H_max_ and M_threshold_, the key parameters for C-EPOCS. Additionally, we demonstrate the feasibility of using an automated response delineation algorithm under these conditions. Despite reduced signal clarity, the algorithm reliably identifies M-wave and H-reflex responses in real time.

Overall, this study demonstrates the feasibility of a portable C-EPOCS system capable of providing immediate feedback based on both EMG and EEG signals. It also offers practical recommendations for selecting acquisition hardware to support reliable signal quality, real-time processing, and portability.

## Introduction

A real time feedback of single trial evoked responses can be useful in developing novel neurorehabilitation applications that target specific neural or spinal evoked responses such as sensory-evoked potentials (SEPs) (Gupta et al., 2025b), spinal reflex responses (H-reflex) (Wolpaw 1987, Thompson and Wolpaw 2021, Thompson et al., 2009, 2013), motor evoked potentials (Thompson et al., 2018), and event related negativity (Chavarriaga et al., 2014, Gomez-Andres et al., 2024, Wirth et al., 2019). Such applications can help enhance weakened responses post injury (Thompson et al., 2018) or suppress maladaptive hyperactive responses (Thompson et al., 2009, 2013), via operant conditioning.

The Evoked Potential Operant Conditioning System (EPOCS) (Hill et al., 2022, Thompson et al., 2009) and Cortical-Evoked Potential Operant Conditioning System (C-EPOCS) (Gupta et al., 2025b) are applications that were developed for such real time closed-loop operant conditioning of evoked responses. These were built as application layers of the BCI2000 software platform (Melinger and Schalk 2007, Schalk et al., 2004), and provide practical interfaces for performing the multiple tasks required for the conditioning.

The EPOCS hardware and software were originally developed to process and provide feedback on evoked responses recorded peripherally using electromyography (EMG) (Thompson et al., 2013; Thompson et al., 2018). In contrast, the C-EPOCS system was customized to process and deliver feedback based on cortical responses recorded via electroencephalography (EEG) (Gupta et al., 2025a). Both systems– particularly C-EPOCS– are component-heavy and have primarily been evaluated in controlled laboratory environments.

In long-term interventional studies, in individuals with brain or spinal cord injuries, these setups may reach fewer people, limiting efficacy testing. Hence, developing a portable and practical version of the system could enable a wider reach, including research in constrained settings, such as rehabilitation centers, and home-based interventions and assessments. A portable system would also allow enhancing the efficacy of these conditioning approaches by combining them with active tasks (Thompson and Wolpaw 2021). All of these would facilitate clinical translation. In addition, the ability to collect longitudinal data from a larger number of people with injuries would also allow mechanistic studies on long-term neuroplasticity.

The C-EPOCS is designed for real-time acquisition and feedback of single-trial cortical evoked responses, typically known to have a low signal-to-noise (SNR) ratio, partly due to the variability in brain responses, but also due to variabilities in the stimulation procedures and stimuli. Focusing on an electrical stimulation-based C-EPOCS, the afferent excitation of nerves can differ across sessions, even with the same magnitude of applied stimulation current, due to small differences in surface electrode positions, posture, and muscle contractions. This makes it challenging to track functional improvements across multi-session interventions. In our previous study (Gupta et al., 2025a), we discuss an approach to maintain the *effective* afferent excitation– i.e., the stimulation received at the afferent pathway– as a means to reduce some of the inter-session variability. We perform this procedure on each testing day by measuring the stimulation intensity– referred to as the *target* stimulation intensity– required to elicit a near-maximal spinal reflex response (Hoffman reflex or H-reflex (Palmieri et al., 2004)), and a 10-20% direct muscle action potential (M-wave), in a muscle innervated by the stimulated nerve. These values are determined by generating recruitment curves for both the H-reflex and M-wave.

In a C-EPOCS study, this is achieved in two steps: first, an instantiation of the standard EPOCS setup (Hill et al., 2022) (Figure 1) is used, to record H-reflex and M-wave recruitment curves. These are used to determine the optimal *target* stimulation intensity for eliciting robust cortical responses (Gupta et al., 2025 a, b). In this setup, the EPOCS computing station receiving the EMG data acts as the control unit and is programmed to trigger the stimulator via a digitizer. Subsequently, the system is switched to the C-EPOCS setup (Figure 2) to record, process and provide feedback on cortical responses at the *target* stimulation intensity. In this setup, the C-EPOCS system receiving the EEG data acts as the control and triggers the stimulator via a programmable embedded system. In these 2 setups, as the EEG and EMG signals are being recorded by separate hardware systems, the EPOCS and C-EPOCS are maintained on independent computing stations to avoid errors during switching of connections.

**Figure 1.**
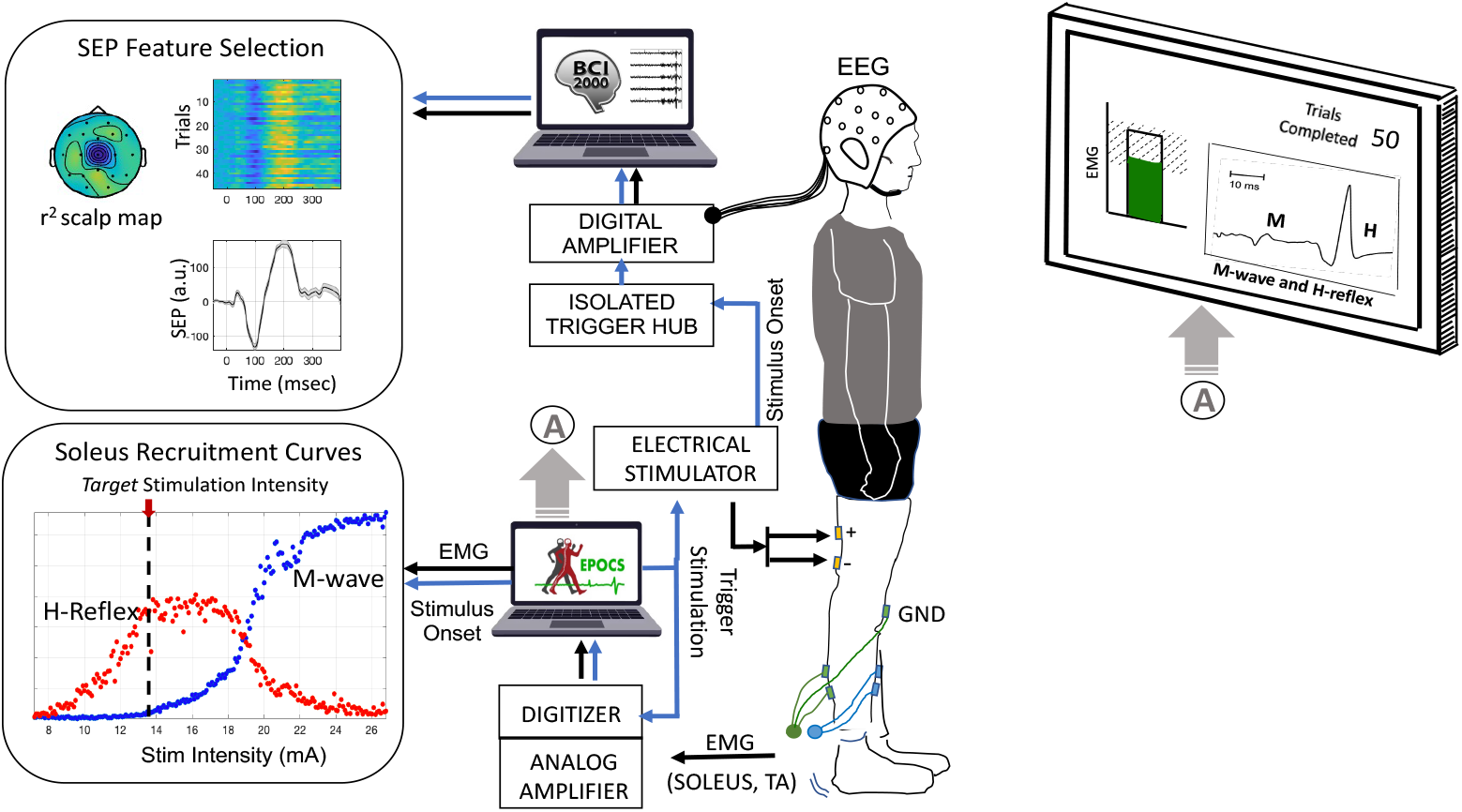
The Evoked Potential Operant Conditioning System (EPOCS) is designed to measure effective afferent stimulation by recording spinal reflex responses—specifically the H-reflex and M-waves—at the soleus muscle during tibial nerve stimulation. The setup includes preamplifiers, an analog amplifier, a digitizer, and an electrical stimulator, forming a well-established framework that integrates with EPOCS software to process incoming electromyography (EMG) data and display real-time responses on a monitor. Data is typically recorded at a high sampling rate of 3200 Hz. EPOCS also continuously monitors background EMG activity in the soleus and its antagonist, the tibialis anterior, ensuring both remain below predefined thresholds before triggering stimulation. Brain signals are recorded separately using electroencephalography (EEG) via an independent biosignal acquisition system running the BCI2000 software platform. EEG and EMG data streams are synchronized by recording a copy of the stimulation trigger onset alongside the EEG data.

**Figure 2.**
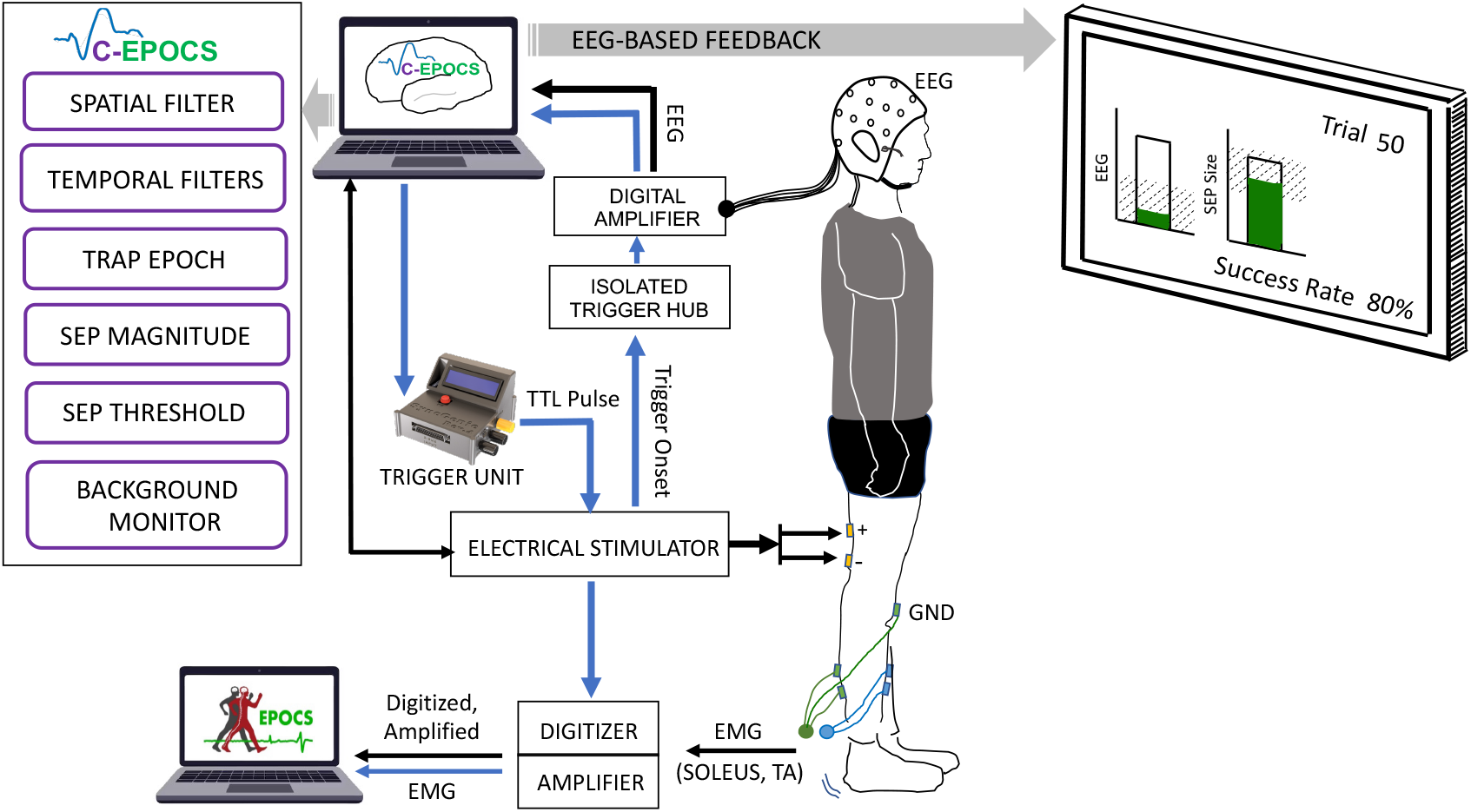
C-EPOCS Setup: This setup primarily records and displays brain responses to electrical stimulation using electroencephalography (EEG). It verifies that the ongoing EEG remains within predefined resting-state limits before triggering stimulation via a programmable Trigger Unit. EEG signals are recorded and processed in real time at 300 Hz, with the resulting sensory responses displayed on a monitor. In parallel, spinal reflex responses (H-reflex and M-wave) are recorded using electromyography (EMG) through a separate standard EPOCS system. EMG is sampled at 3200 Hz using an acquisition system (amplifier and digitizer) integrated with EPOCS. A copy of the stimulation trigger is recorded within the EPOCS system to synchronize the EEG and EMG data streams. [Adapted version of Figure 2–Gupta et al., 2025b]

This overall system includes several bulky hardware components that require careful assembly and handling by trained researchers. These include EEG electrode setup with pre-amplifiers, amplifiers, digitizers, trigger hubs; EMG analog amplifiers with preamplifier unit, battery unit, digitizer unit, electrical stimulation device; two computing stations with extended viewing monitors for participant and researcher; synchronization embedded system; multiple power sources and various connector cables for communication and synchronization of these hardware systems. The setup becomes expensive and requires planning and organization on specialized carts, with several potential points of failure. Some of these components–such as the analog amplifier with a sampling rate of 3200 Hz–may not be readily available in many laboratories and the whole setup can be burdensome to setup in space constrained research or clinical settings.

In this study, we present the customizations and evaluations performed to streamline the above two steps (EPOCS and C-EPOCS) into a single, compact, user-friendly platform that eliminates the need for manual component switching. The system is designed to acquire and process both cortical and muscle responses, support contingent response processing, trigger electrical stimulation, and deliver rapid, real-time feedback on muscle (H-reflex, M-wave) or cortical activity through visual gauges, with a single system.

## Methods

A key feature of the portable setup is the use of a single biosignal recording device— such as a dry, active, battery-powered EEG headset (with optional wireless capability)—that can simultaneously acquire EMG signals and includes an embedded, isolated port for digital triggers. Important criteria in selecting the data acquisition hardware include high common-mode rejection ratio, integrated pre-amplifiers in EEG/EMG electrodes, high sampling rate, and compatibility with platforms like BCI2000. We performed the assessment with a DSI-24 system (Wearable Sensing, CA), but other EEG acquisition systems with similar capabilities may also be suitable. The main limitation of recording EMG with an EEG-based system (especially in a portable or wireless system), is the lower sampling rate, which affects the EMG signal resolution.

We evaluate whether the reduced EMG signal resolution impacts the measurement of M-wave and H-reflex responses, and whether their recruitment curves can still be generated in real time to obtain target stimulation intensity accurately, without requiring extensive post-processing. To address this, we first conduct a pseudo-online analysis in which EMG signals were acquired at a high sampling rate (3200 Hz). The stimulation intensities that elicited maximal H-reflex (H_max_) and M-wave (M_max_) responses were then compared with those obtained from down sampled, lower-resolution EMG signals—specifically, at 640 Hz (5× lower) and 320 Hz (10× lower). These sampling rates were selected as they closely match the sampling rates typically used for EEG systems. The subjective delineation of these responses can become challenging at reduced EMG resolution and an automated approach can be useful. Towards that, we test the performance of an automated delineation algorithm (McKinnon et al., 2023) designed for H-reflex studies, on these lower resolution EMG signals. Next, we test the portable system in an online mode to evaluate its operational feasibility as it simultaneously records and transmits EEG and EMG signals at 600 Hz (a pre-defined setting) to the C-EPOCS system. We assess the resulting M-wave and H-reflex recruitment curves, by curve fitting and quantifying curve parameters. We focus in particular on the stimulation intensities required to reach H_max_ and M_max_, as they are used for obtaining the target stimulation intensities for use in the C-EPOCS.

### Portable C-EPOCS architecture

In the standard setup (illustrated in Figure 1), EMG is recorded at the soleus muscle, using an analog amplifier (AMT-8, Bortec Biomedical Ltd., Canada) at a sampling rate of 3200 Hz. The system includes a battery-powered belt unit with bipolar electrodes and preamplifiers, connected to an analog-to-digital converter (PCIe-6321, National Instruments, TX). The digitizer is linked to a computing station running the EPOCS software platform, which serves as the control unit. This control station continuously receives and processes EMG signals to assess whether the background EMG activity in the soleus muscle (or antagonist tibialis anterior muscle) is maintained at a predefined low level (approximately 5– 10% of maximum voluntary contraction or 5-18 *µ*V observed in soleus during normal standing) for at least 2 seconds. Upon meeting this condition, the system triggers the electrical stimulator according to the selected protocol—for example, in recruitment curve protocol, it systematically increases stimulation intensity in predefined steps after a set number of trials. The control station also receives stimulation onset timestamps and other trial-specific parameters directly from the stimulator. Prior to each trial, the background EMG is continuously monitored, processed, and displayed on a participant-facing monitor to allow them to maintain it at the required low level. This setup is used to acquire the recruitment curves of the H-reflex and M-wave, enabling selection of a target stimulation intensity at which the H-reflex amplitude is approximately 75% of H_max_ and the M-wave amplitude is around 10% of M_max_. These procedures have been described in Hill et al., (2022) and used in multiple studies (Thompson et al., 2021; Gupta et al., 2025a). In this setup, the brain signals are recorded simultaneously separately with an EEG acquisition system (including pre-amplifier, amplifier, digitizer, and trigger hub). The EEG and EMG data streams are merged by synchronization pulses from the electrical stimulator.

After determining the target stimulation intensity with the help of the standard EPOCS, the setup it switched to C-EPOCS to provide feedback based on brain signals. Switching to C-EPOCS involves transferring control from the standard EPOCS system (Figure 1)—that processes EMG signals and triggers the stimulator via a digitizer—to the C-EPOCS system (Figure 2), that processes EEG signals and triggers the stimulator via the synchronization unit. Since the wiring configurations for the two systems differ, they are maintained on separate computing stations to prevent connectivity errors during the rapid switch-over. The C-EPOCS system communicates with the synchronization embedded system (SyncGenie, Alkhoury et al., 2024) which in turn transmits a digital trigger to the electrical stimulator. Stimulation parameters for each trial are collected directly from the stimulator. EEG is recorded using a 19-channel referential dry electrode headset (DSI-24, Wearable Sensing) at a sampling rate of 300 Hz, with data acquired via the BCI2000 software platform. The EEG electrodes include built-in preamplifiers, and the headset is battery-powered. C-EPOCS also retrieves key EEG metadata from the headset, such as channel configuration, sampling rate, and timestamps.

The proposed portable setup (illustrated in Figure 3) consists of only four components: (a) the EEG headset (DSI-24, Wearable Sensing), for simultaneously recording of synchronized EEG and EMG signals; (b) the electrical stimulator (DS8R, Digitimer), for delivering electrical stimuli; (c) the synchronization embedded system (SyncGenie, Alkhoury et al., 2024), for synchronizing the data streams; and (d) a computing station (laptop or desktop) with an extended monitor, to operate the C-EPOCS and display feedback. In this setup, the EEG and EMG data are both acquired at a sampling rate of 600 Hz, a typical rate supported by the EEG headset. EMG is recorded from one of the DSI-24’s three auxiliary channel inputs, synchronously digitized by the same ADC electronics as the EEG. Both EEG and EMG are active electrodes with built-in pre-amplifiers. All biosignals and trigger recordings are transmitted to the C-EPOCS system with precise time synchronization.

**Figure 3.**
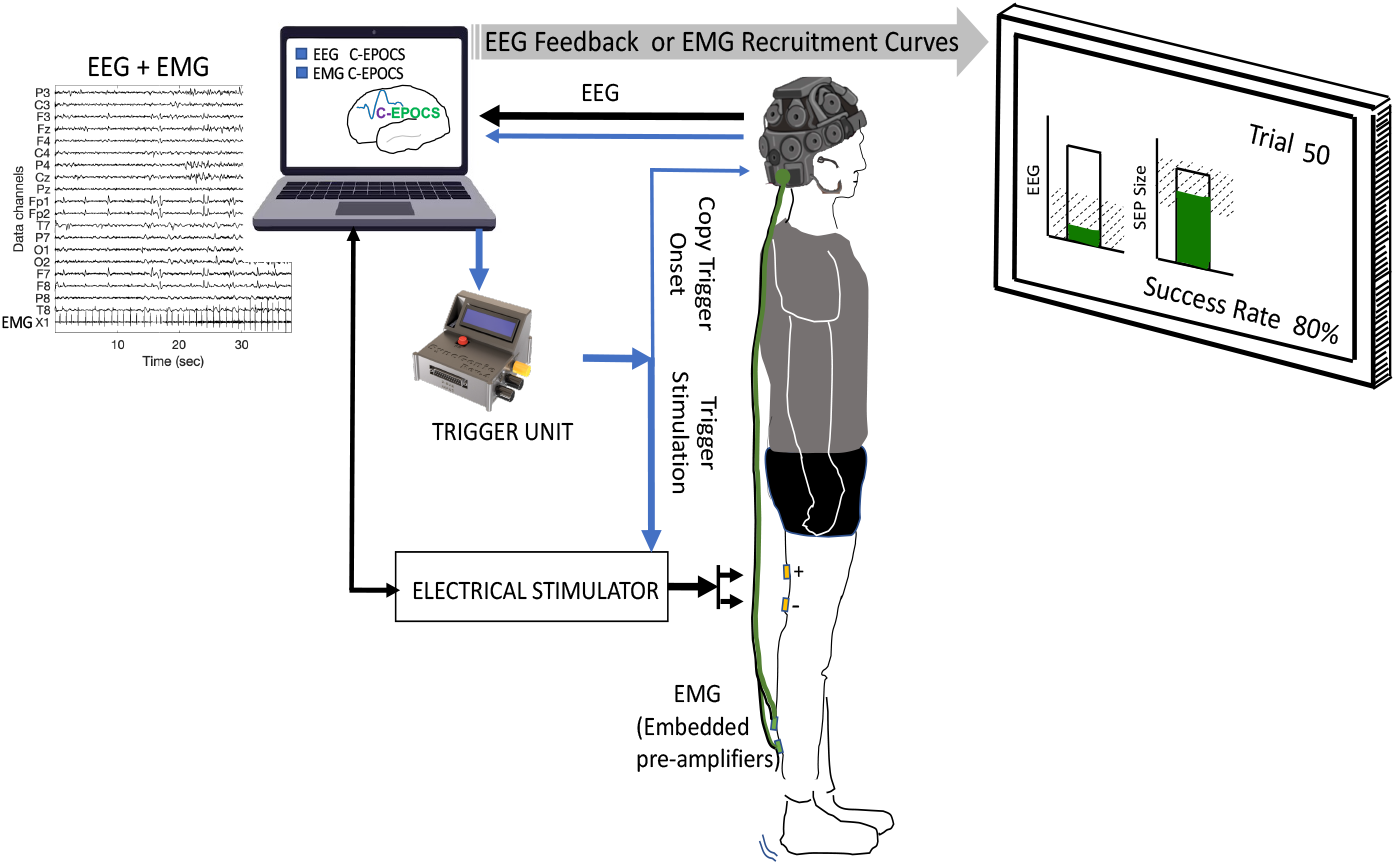
Portable C-EPOCS Setup: This configuration records both brain signals (electroencephalography, EEG) and muscle signals (electromyography, EMG) at 600 Hz, using a single acquisition system—comprising the EEG headset and C-EPOCS software. Designed with a smaller footprint, the system can seamlessly switch between standard EPOCS functions for EMG-based feedback and C-EPOCS functionality for EEG-based feedback—without requiring any changes to the wiring or equipment setup. In EMG mode, C-EPOCS monitors background EMG activity prior to triggering stimulation and displays the resulting recruitment curves. In EEG mode, it monitors resting-state EEG before stimulation and displays sensory evoked potentials. A programmable Trigger Unit transmits digital pulses to the stimulator to initiate each trial.

The C-EPOCS is customized with an additional skin (INI settings configuration) to easily switch between an EMG-driven system to an EEG-driven system, with the hardware and cable connections remaining unchanged. This setup integrates the functionalities of standard EPOCS and C-EPOCS setups and can be operated using a single standard laptop or desktop computer. This development simplifies the setup and reduces the hardware footprint and potential points of failure in switching between EPOCS and C-EPOCS systems.

### Portable C-EPOCS evaluation

We evaluate the portable C-EPOCS setup in two steps: i) a pseudo-online analysis that used a stand-alone standard EPOCS for obtaining recruitment curves, and ii) testing the system’s operational feasibility in an online mode where the EEG and EMG were recorded simultaneously and processed through a single EPOCS system.

### Data

Retrospective analysis was performed on data from 21 healthy participants (age range: 23-82 years; mean age: 48.3y + 18.8y, 10 women and 11 men), with no known neurological injury, and one individual with chronic stroke (> 18-years-old, stroke more than 6 months ago; age and gender are not disclosed to maintain de-identification). The data was collected during methods development study on H-reflex and somatosensory evoked potentials (Brangaccio et al., 2025), and baseline measurements during a motor learning study (Mojtabavi et al., 2025, Gupta et al., 2025c), at our laboratories. Real time assessment was performed in 4 participants from the healthy participant group (36y + 14.9y, 2 women and 2 men). All data was acquired with informed consent, and the protocol (# 1584762) was approved by the local institutional review board at the Stratton Veterans Affairs Medical Center.

Both the datasets were obtained during electrical stimulation of the tibial nerve at the popliteal fossa. The stimulation was delivered at 1 Hz stimulation frequency with an inter stimulus interval (ISI) jitter of 10%, with 1 msec long biphasic pulses. Stimulation was delivered with a constant current stimulator (DS8R, Digitimer) with surface electrodes, with the cathode (proximal) and anode placed at 3 - 4 cm inter-electrode distance. The stimulation and recording setup are as described in previous studies (Hill et al., 2022, Gupta et al., 2025a, Thompson et al., 2013). To obtain the H-reflex and the M-wave recruitment curves, the stimulation current was increased steadily from threshold (minimum stimulation current that elicits an H-reflex) till a maximal M-wave was observed, which asymptotes with further increase in stimulation intensity.

### EMG recording

In the standard setup, EMG was recorded using a pair of bipolar electrodes with built-in preamplifiers (500x gain) (Bortec Biomedical Ltd.) at 3200 Hz sampling rate. In the portable setup, we used bipolar EMG electrodes (EXG-Bi-60, Wearable Sensing) with integrated preamplifiers (60x gain), connected to auxiliary channels in the EEG headset (DSI-24, Wearable Sensing), sampled at 600 Hz, analog high pass filtered at 0.003 Hz and digitized by the same ADC as the EEG Electrodes. Electrodes were attached to the skin using self-adhesive Ag/AgCl surface electrodes (Neuroplus, 22 × 22 mm, A10040-60, Vermont Medical Inc.) placed over the soleus muscle belly, approximately 3–4 cm apart (center-to-center).

### EMG analysis

In the retrospective analysis, EMG data were low-pass filtered at 200 Hz (2nd-order Butterworth filter) and then down sampled to 640 Hz (5× lower than the original 3200 Hz), or low-pass filtered at 100 Hz (2nd-order Butterworth filter) and down sampled to 320 Hz (10× lower). The preprocessed data were then segmented into 600 msec epochs (from −200 to 400 msec relative to stimulation onset). Next, the H-reflex and M-wave response time windows were determined in two ways: (a) subjectively by the investigator in consultation with an expert in the laboratory, and (b) objectively using an algorithm specifically designed for automated delineation of these responses (McKinnon et al., 2023). Subsequently, the mean rectified values (MRV) of the signal within these time windows were calculated. Next, the recruitment curves were obtained at the three sampling rates (3200 Hz, 640 Hz, 320 Hz) by plotting these MRV values at the respective stimulation intensities. Finally, to compare curve parameters across sampling rates, the H-reflex and M-wave recruitment curves were fitted with a bell shaped and a sigmoid curve respectively, as described in McKinnon et al., (2023) using the open-source software shared by the authors. The automated delineation can help to reduce variability that arises from subjective evaluation, especially in longitudinal studies, but it may fail at the lower resolution EMG signals. The process includes blanking the stimulus artifact; removing the stimulus artifact related decay with an exponential function; modelling the M-wave as a logistic function of stimulus intensity and modelling the H-reflex with the ‘hill curve’– a product of two sigmoids. Six curve parameters were extracted for comparison–residual errors for the H-reflex and M-wave curves quantified by the mean squared error (MSE), H_max_, M_max_, stimulation intensity at H_max_ and stimulation intensity at M_threshold_. These parameters were selected as they are most relevant for the C-EPOCS protocol, where the stimulation intensity of H_max_ and M_threshold_ is determined as the target stimulus intensity for maintaining the afferent stimulation.

In the real time demonstration of portable C-EPOCS, the EMG was filtered with a 10 Hz high pass filter (Butterworth, model order 2), to remove movement related artifacts, generally observed in data acquired by the headset at the higher stimulation intensities. The subsequent procedure for recruitment curve fitting remained the same as the retrospective analysis.

### EEG recording and analysis

In both the setups (retrospective and portable), 19 channel EEG was recorded with the dry active DSI-24 headset, using referential mode. The electrodes were pre-positioned according to the International 10-20 EEG system, with the ground at Fpz and the reference at linked earlobes. In the retrospective dataset, the EEG was collected at 300 Hz, while in the portable C-EPOCS setup, the EEG was recorded at 600 Hz along with the EMG. The EEG data is not expected to be affected in the transition from the standard to the portable setup, hence these are not included in the current analysis.

### Statistical Analysis

All analysis was performed in MATLAB 2020b (Mathworks, MA). Repeated measures assessments across sampling rates tested in the same group of participants were performed with Friedman Repeated measures test, followed by *post-hoc* tests with Tukey’s multiple comparison procedure.

### Results

We evaluate the effect of low-resolution (low sampling rate) EMG signals on the H-reflex and M-wave recruitment curve parameters, especially the stimulation intensity at H_max_ and M_threshold_, important for C-EPOCS. First, we analyze existing data in healthy people and the individual with stroke, by down sampling the EMG signals from 3200 Hz to 5x lower (640 Hz) and 10x lower (320 Hz) sampling rate, identifying the H-reflex and M-wave response time windows by hand, fitting their recruitment curves at each sampling rate, and comparing the curve parameters. We hypothesized that curve parameters may be affected as the EMG resolution is reduced. We also assess the above with the automated delineation of responses as described in McKinnon et al., (2023). We hypothesized that the automated response delineation algorithm will not be able to perform as well at the lower resolution EMG. Lastly, we assess the operational feasibility of the portable C-EPOCS, i.e., recording both EEG and EMG with the EEG headset at its typical sampling rate (600 Hz), processed in real time via the portable C-EPOCS pipeline.

EMG data from 21 healthy participants (age: 23-82 years) and 1 participant (> 18 years old) with neurological injury was analyzed. Data was recorded at the soleus muscle during tibial nerve stimulation. In all participants, for the recordings at 3200 Hz, we observed the expected recruitment curves for H-reflex (bell-shaped curve) and M-wave (sigmoid curve) (shown by a representative example in Figure 4).

**Figure 4.**
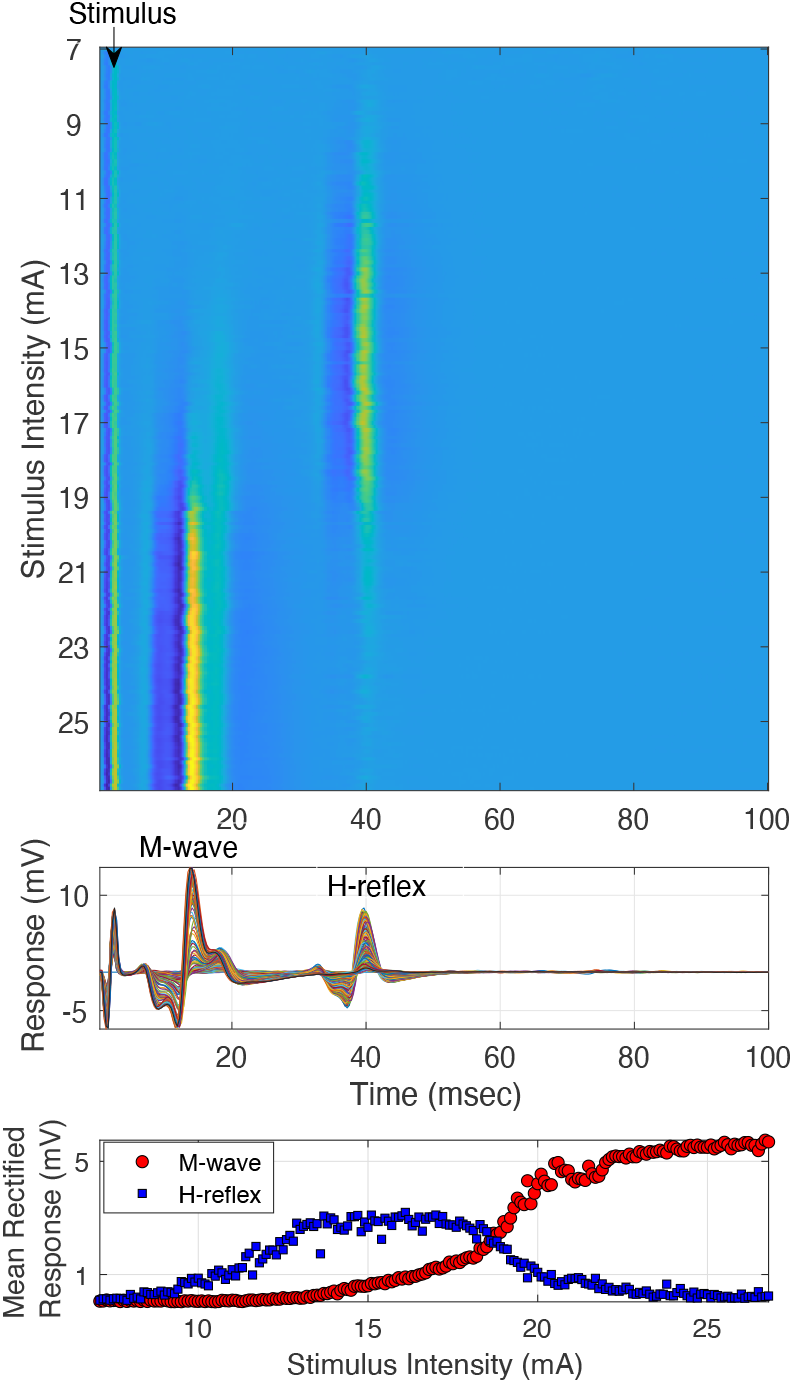
Representative Example: M-wave and H-reflex recruitment curve data obtained at 3200 Hz sampling rate, recorded at the soleus muscle, during tibial nerve stimulation at the popliteal fossa, in a healthy participant. Top panel: Heat plot of all the trials, each row being a trial, and color the EMG response magnitude. Middle panel: Trials collapsed as overlapped time courses, show the M-wave and H-reflex responses. Lower panel: recruitment curve data plotted as the mean rectified M-wave and H-reflex responses at each stimulation intensity.

### Pseudo-online assessment with down sampled EMG recordings in healthy people

The recruitment curves fitted with EMG at 3200 Hz, 640 Hz and 320 Hz visually showed considerable difference and deterioration with the 10x lower sampling rate, and lesser change with 5x lower sampling rate. The M-wave curves appeared to be more affected while the H-reflex curves appeared more robust. A representative example shown in Figure 5 illustrates the increased noise in the M-wave as the sampling rate was reduced from 3200 Hz to 320 Hz. This change was quantified by the mean squared error of the residuals. Despite this increased dispersion of data points, the curves did not appear to shift, quantified by comparing the stimulation intensity at H_max_ and M_threshold_ at the different sampling rates. We also assess the magnitude of H_max_ and M_max_, which can be affected by the filtering process. The descriptive statistics (mean and standard error) of these curve parameters across participants are shown in Figure 6.

**Figure 5.**
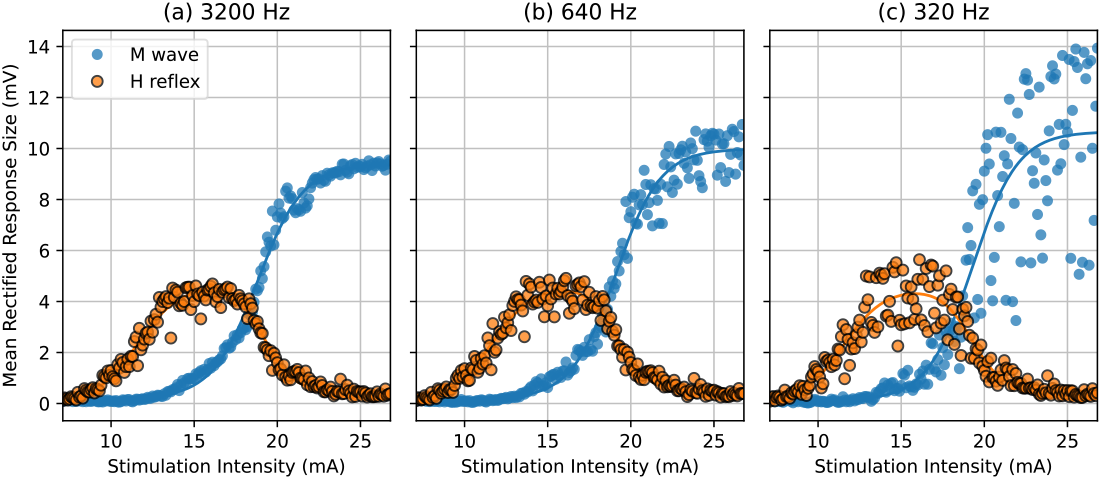
Representative recruitment curves from one healthy participant. EMG recorded at (a) 3200 Hz, (b) 640 Hz and (c) 320 Hz. The lower sampling rate shows increased residuals for both H-reflex and M-wave data, especially at higher stimulation intensities.

**Figure 6.**
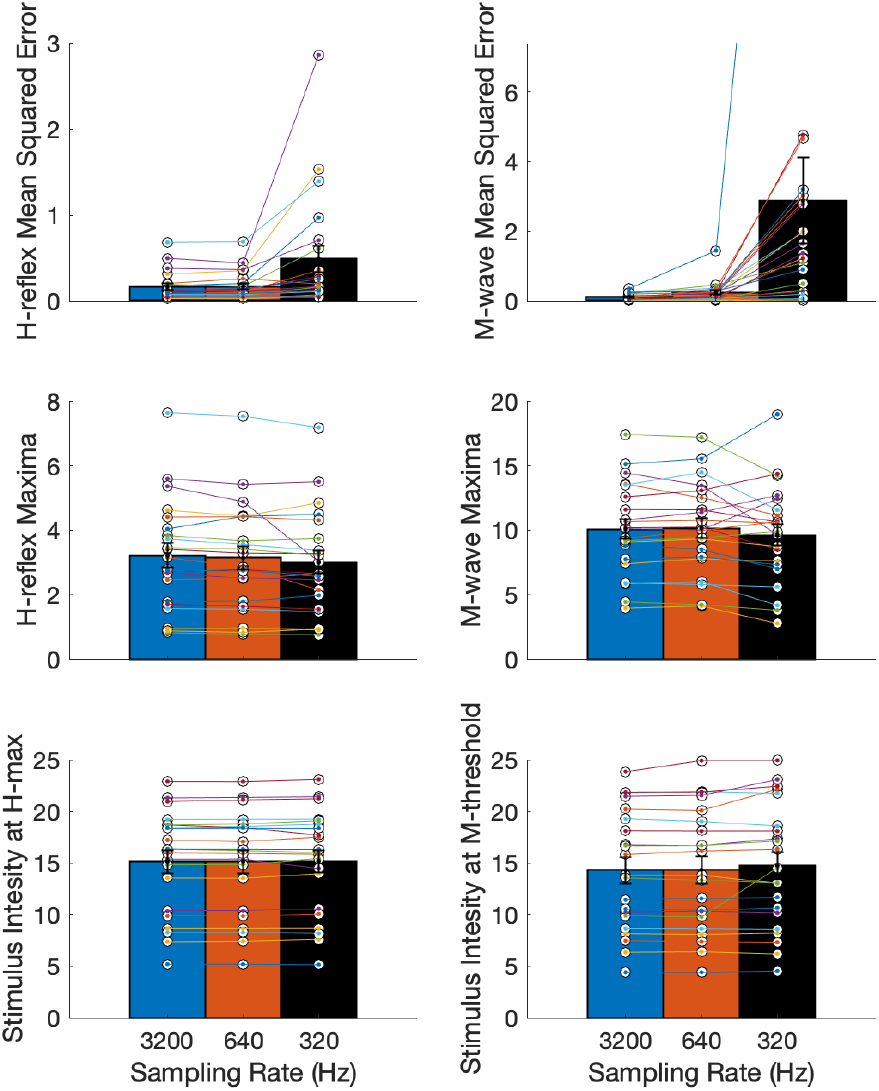
Curve parameters with investigator selected response windows for H-reflex and M-wave at the three sampling rates (3200 Hz, 640 Hz and 320 Hz) (mean, standard error). Top panel: Mean squared error (MSE) for H-reflex and M-wave recruitment curves showed increased MSE at lower stimulation intensity (Friedman repeated measures test, p < 0.001 for both). Middle panel: Magnitude of H-reflex and M-wave at their recruitment curve maxima; maxima decreased with lowering of sampling rate (p < 0.05 for H-reflex). Bottom panel: Stimulation intensity at H_max_ and M_threshold_ remained unchanged with lower sampling rate (p > 0.05 for both).

The Friedman repeated measures test showed a significant difference between the residual errors or MSE for H-reflex and M-wave at the three sampling rates (χ^2^ = 20.67, *p* = 3.253*e*^-9^) for H-reflex, and (χ^2^ = 40.09, *p* = 1.965*e*^-9^) for M-wave. The *post hoc* test showed a significance difference between H-reflex at 3200 Hz and 320 Hz (*p* = 0.0002) and between 640 and 320 Hz (*p* = 0.0003), and a significant difference in M-wave between all pairs of sampling rates (*p* < 0.01). The H_max_ was observed to be significantly different across sampling rates as well (χ^2^ = 8.67, *p* = 0.013); with the *post hoc* test indicating significant difference between 3200 Hz and 320 Hz (*p* = 0.009). The M_max_ was not found to be significantly different (χ^2^ = 5.43, *p* = 0.066).

However, more importantly, the stimulus intensity at H_max_ and M_threshold_ were not found to be significantly different across the three sampling rates (χ^2^ = 1.81, *p* = 0.405) and χ^2^ = 1.24, *p* = 0.538). This is useful for C-EPOCS application, where the stimulus intensity at H_max_ and M_threshold_ are the main parameters of interest from these recruitment curves.

### Automated response delineation and recruitment curve fitting

Next, we assessed the use of an automated process to delineate the H-reflex and M-wave responses (McKinnon et al., 2023), which we hypothesized to be considerably challenging for signals with a lower resolution. We fit the curves as described in the previous sections and compare the curve parameters: MSE for H-reflex and M-wave, magnitude of H_max_ and M_max_, and stimulation intensity at H_max_ and H_threshold_, across the three sampling rates. The descriptive statistics (mean and standard error) of these curve parameters across participants are shown in Figure 7.

**Figure 7.**
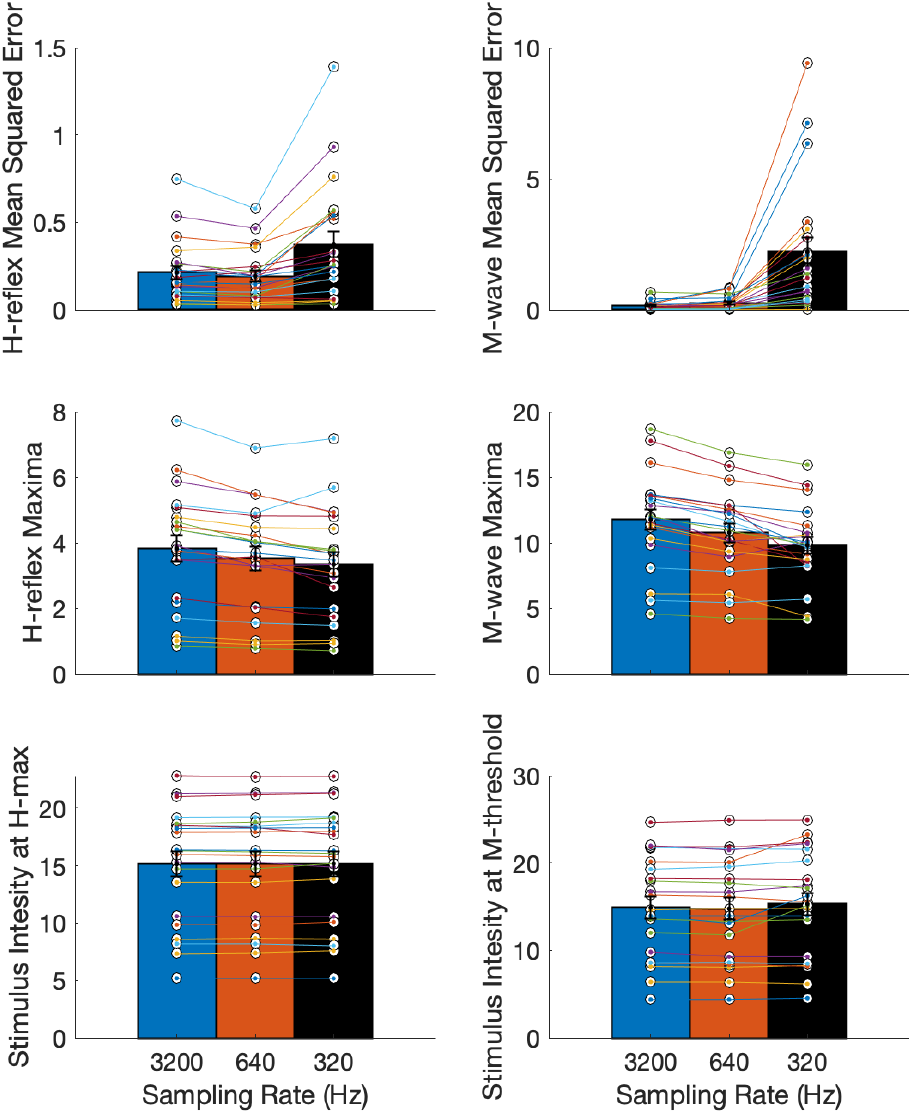
Curve parameters with automated delineation of H-reflex and M-wave responses at the three sampling rates (3200 Hz, 640 Hz and 320 Hz) (mean, standard error): Top panel: Mean squared error (MSE) for H-reflex and M-wave recruitment curves showed increased MSE at lower stimulation intensity (Friedman repeated measures test, p < 0.001 for both). Middle panel: Magnitude of H-reflex and M-wave at their recruitment curve maxima decreased with lowering of sampling rate (p < 0.001 for both). Bottom panel: Stimulation intensity at the H_max_ and M_threshold_ remained unchanged with lower sampling rate (p > 0.05 for both).

The Friedman repeated measures test showed that the residuals of H-reflex and M-wave recruitment curves (quantified by the mean squared error (MSE)) were significantly different across sampling rates (χ^2^ = 27.81, *p* = 9.15*e*^-7^ for H-MSE and χ^2^ = 35.53, *p* = 1.93*e*^-8^ for M-MSE). *Post hoc* tests showed significant difference in H-MSE and M-MSE between 3200 Hz and 320 Hz (*p* < 0.01), and between 640 Hz and 320 Hz (*p* < 0.01). The H_max_ and M_max_ magnitudes showed a significant difference with sampling rate (χ^2^ = 28.67, *p* = 5.96*e*^-07^ and χ^2^ = 24.67, *p* = 4.40*e*^-6^) respectively. *Post hoc* tests showed significant difference between 3200 Hz and 640 Hz (*p* < 0.01), and between 3200 Hz and 320 Hz (*p* < 0.01) for both H_max_ and M_max._ However, the stimulation intensity at H_max_ and M_threshold_ did not differ significantly (χ^2^ = 1.52, *p* = 0.47 and χ^2^ = 3.52, *p* = 0.17, respectively), which is useful as it indicates that this parameter is robust to lower resolution EMG.

### Pseudo-online assessment in individual with neurological injury

The retrospective analysis on the data from the individual with neurological injury showed typical H-reflex and M-wave recruitment curves (Figure 8). We observe a small change in the H-reflex residuals at lower sampling rates (6.1% at 640 Hz and 22.9% at 320 Hz) and a relatively larger change in the M-wave residuals (13.3% at 640 Hz and 76.92% at 320 Hz). A small decrease was observed in the H_max_ (-5.1% at 640 Hz and -12.9% at 320 Hz) and M_max_ (0.1% at 640 Hz and -4.1% at 320 Hz) magnitudes. The stimulation intensities at H_max_ and M_threshold_ had a negligible change–0.4% at 640 Hz and 1.5% at 320 Hz for H_max_ and 0.6 at 640 Hz and 1.6% at 320 Hz for M_threshold_. These trends were similar to that observed in the participants with no neurological injury.

**Figure 8.**
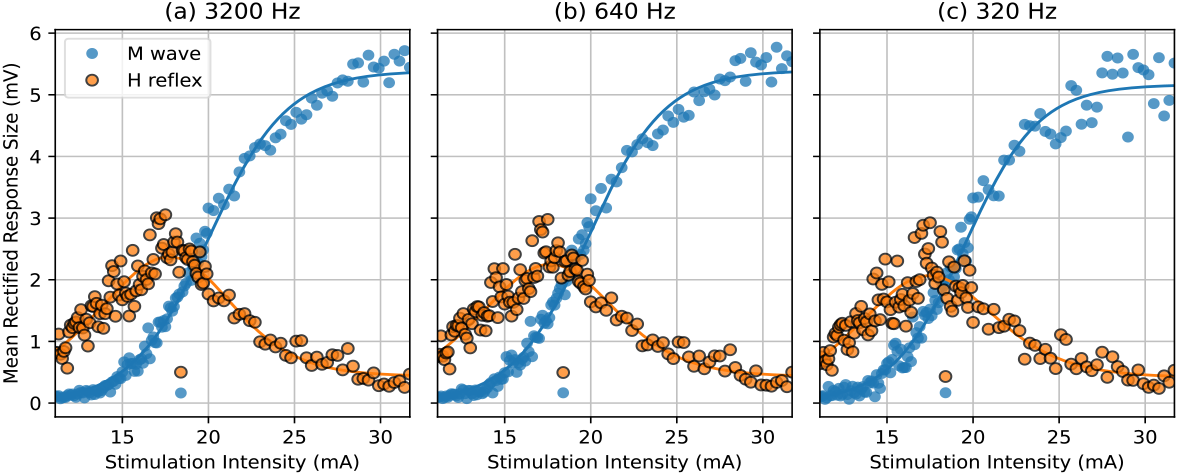
Recruitment curves for H-reflex and M-wave measured in an individual with stroke: The M-wave residuals show some dispersion at 320 Hz at higher stimulation intensities, but mainly show that the curves do not change drastically with the lower sampling rate, such as at 640 Hz. The stimulation intensity at H_max_ shows a negligible change of 0.4% and 1.5% and at M_threshold_ 0.6% and 1.6% at 640 Hz and 320 Hz respectively.

### Demonstration of real time portable C-EPOCS system

Here, we demonstrate the operational feasibility of recording the H and M recruitment curves with the portable setup (illustrated in Figure 3). Here the EMG is recorded along with the EEG using a common hardware device– the EEG headset. Both the datasets are recorded at 600 Hz. This data was collected in 4 participants (age: 23-57 years, 2 women and 2 men) with no known neurological injury. The recruitment curves were observed to be the typical shape similar to the retrospective analysis dataset. A representative example is shown in Figure 9. The residuals (quantified by mean squared error) for the H and M recruitment curves were assessed as a measure of dispersion of points around the curve. The MSEs for H-reflex and M-wave curves were negligible – 0.014 + 0.005 and 0.010 + 0.001 respectively, when analyzed with the investigator selected response windows; and 0.018 + 0.006 and 0.013 + 0.003 respectively, with the automated response delineation.

**Figure 9.**
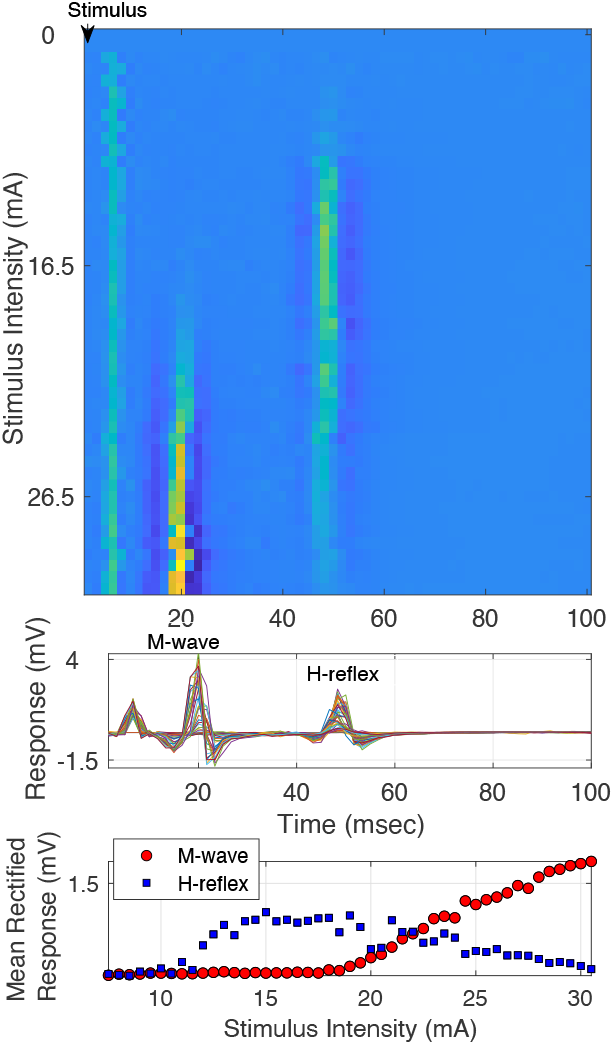
Measuring H-reflex and M-wave recruitment curves with the EMG acquired with the EEG headset, at the lower sampling rate. Top panel: Heat plot of all the trials across time. The color represents the magnitude of the signal. Middle panel: Shows time courses of the trials in a butterfly plot. The M-wave and H-reflex can be observed clearly despite lower resolution. Bottom panel: Recruitment curves: Averaged H-reflex and M-wave responses plotted across stimulation intensities.

## Discussion

The C-EPOCS system is designed to provide real-time feedback based on cortical responses to individual trials. However, these single-trial cortical signals typically have a low signal-to-noise ratio (SNR) and are highly susceptible to artifacts and background noise. This poses challenges for real-time applications, where feedback must be timely and accurate without extensive averaging or post-processing.

In conventional protocols, the intensity of electrical stimulation is typically determined using subjective measures, such as visible muscle twitches or reported sensory perception. However, the nerve excitation can vary across sessions due to changes in electrode positioning, participant posture, muscle tone, or amount of involuntary contraction—factors that are especially difficult to control in individuals with brain or spinal cord injuries. Therefore, despite maintaining a constant stimulation current, the actual stimulation received by the nerves can differ across sessions, introducing variability in afferent input and affecting the ability to track intervention-related cortical changes over time.

To address this, it can be useful to objectively assess and standardize afferent excitation across sessions. Inspired by established H-reflex conditioning protocols, our previous work (Gupta et al. 2025a,b) described a procedure that uses H-reflex and M-wave recruitment curves to identify a target stimulation intensity that elicits consistent single trial cortical responses. However, these methods were developed using high-resolution EMG recordings sampled at 3200 Hz, requiring a component-heavy setup that limits portability and usability in non-laboratory environments.

In this study, we explored whether the same objective procedures could be adapted for a more portable system—one that combines EEG and EMG data acquisition into a single device, typical of mobile neurotechnology platforms. A key trade-off in such systems is the reduced sampling rate, which can be 5–10 times lower than that used in traditional high-fidelity setups.

Despite this limitation, our results show that H-reflex and M-wave recruitment curves can still be generated using low-resolution EMG signals, enabling the objective determination of target stimulation intensity even in a portable setting. This makes it possible to standardize afferent input across sessions, supporting more stable cortical responses and improving the interpretability of intervention outcomes.

### Feasibility of maintaining effective afferent excitation with low-resolution EMG

Results indicate that EMG signals sampled at rates five times lower than the standard 3200 Hz (i.e., ∼600–640 Hz) can still be used effectively to determine target stimulation intensity—the key parameter for maintaining consistent afferent excitation across sessions. Under these conditions, the stimulation intensity required to evoke comparable reflex responses remained largely unchanged, supporting the feasibility of using lower-resolution EMG for this specific application.

However, we also observed that further reductions in sampling rate—such as 10x lower (∼320 Hz)—led to increased dispersion in responses at higher stimulation intensities, making the recruitment curves less reliable. Moreover, quantitative parameters such as H-reflex and M-wave magnitudes were notably affected at lower sampling rates, likely due to loss of waveform fidelity. These distortions introduce challenges in applications where precise measurement of response amplitude or latency is required. In such cases, low-resolution EMG should be avoided, and higher sampling rates are recommended to preserve signal accuracy.

We attribute the increased residuals and reduced response amplitudes primarily to the loss of temporal resolution in measuring the stimulus trigger onset, caused by digitization limitations at lower sampling rates. This imprecision results in jitter in the estimated onset of the H-reflex and M-wave, effectively spreading the response time window and reducing the sharpness of the detected waveform.

To mitigate this issue, a potential strategy is to use the stimulus artifact—a large electrical signal that immediately follows stimulation—as a reference for trigger alignment. By estimating the jitter from the variability in the stimulus artifact timing, it may be possible to compensate for timing inaccuracies and improve the reliability of response detection even at lower sampling rates.

### Applicability of Automated response delineation algorithms

Accurate identification of M-wave and H-reflex responses is essential for obtaining the recruitment curves and determining the target stimulation intensity for the C-EPOCS system. However, manual delineation of these responses is subject to inter-rater variability, inconsistent thresholds, and potential inaccuracies, especially when EMG signals are noisy and / or responses are atypical or small. This subjectivity can introduce error into the measurement of key parameters. Algorithm-based delineation methods offer a consistent and objective alternative, reducing the variability introduced by manual marking. Prior work, such as McKinnon et al. (2023), has shown the benefits of automated approaches in improving the reproducibility and robustness of H-reflex and M-wave quantification, as used on high resolution EMG data (recorded at 3200 Hz sampling rate).

In this study, we demonstrate that an automated response delineation algorithm can be successfully applied even when using low-resolution EMG signals (i.e., 5-10x lower sampling rate, such as 640 and 320 Hz)—a common limitation in portable or wireless biosignal acquisition systems. While the clarity of the M-wave and H-reflex responses may deteriorate at lower sampling rates, the algorithm remains robust enough to identify these responses and generate recruitment curves in real time, without the need for extensive post-processing. This finding supports the feasibility of integrating automated response processing into portable C-EPOCS setups, where real-time performance and reduced reliance on expert supervision can be useful. Future work may further improve algorithmic performance by including signal enhancement techniques or addressing trigger jitter issues associated with low-resolution signals.

### Recommended Features for EEG-Based Acquisition in Portable C-EPOCS Systems

The selection of an EEG-based biosignal acquisition system is a critical component in the portable C-EPOCS setup. Below are the key features to consider when evaluating such a system:

a. *Reliable signal acquisition:* The system must reliably capture both cortical and muscle activity with minimal noise and latency, while being compact and user-friendly for out-of-laboratory use.
b. *High sampling rate:* Support for multichannel biosignal recording at a sampling rate of 600 Hz or more is important. This higher temporal resolution will enable accurate capture of fast, transient signals such as M-waves and H-reflexes—key components in a C-EPOCS setup— where standard EEG sampling rates may be insufficient.
c. *Integrated pre-amplifiers:* Pre-amplifiers integrated directly at the electrode level can be useful to reduce motion artifacts and enhance signal fidelity, for single trial responses, especially in ambulatory settings or when working with individuals with injuries, where environmental noise and participant movement are more pronounced.
d. *Consistent electrode placement:* A robust and repeatable method for electrode placement and positioning is important to ensure consistent EEG signal quality across sessions. Variability in placement can significantly affect interpretation of cortical responses, particularly during longitudinal tracking.
e. *EEG-EMG signal isolation:* Isolation between EEG and EMG acquisition pathways is critical. Without it, electrical stimulation artifacts detected by EMG electrodes may contaminate EEG recordings. Hardware-level isolation or signal routing is recommended.
f. *Signal quality and synchronization:* High common mode rejection ratio and high SNR are necessary for both EEG and EMG channels to ensure clean recordings in non-shielded environments. At least one isolated digital I/O port is also essential for receiving stimulation triggers and maintaining precise stimulus-response timing.
g. *Platform compatibility and portability:* Compatibility with real-time platforms such as BCI2000 will enable integrated stimulus control, signal monitoring, and feedback generation. The system should be portable, with amplifiers and digitization components built into the headset to reduce external components, minimize setup time, and increase usability in clinical or home-based settings.

Together, these hardware and software features form the foundation for a robust, portable C-EPOCS system that can support real-time neurofeedback and rehabilitation applications outside traditional lab environments. A more practical and streamlined C-EPOCS setup will also enable integrated research on cortical and spinal responses, expanding its utility for multi-system neurorehabilitation studies.

### Limitations

The retrospective analysis involved the application of low-pass filters prior to down sampling, to prevent aliasing. This preprocessing step can overestimate the effect of reduced sampling rate, depending on the filters used. In contrast, data acquisition systems that are designed to record signals at lower sampling rates often include dedicated hardware and software filters tuned for those rates. These may introduce less distortion or signal attenuation compared to retrospective digital filtering. The real-time data collected at 600 Hz showed smaller residuals than the retrospective analysis of data acquired at a similar sampling rate of 640 Hz.

### Future work

Ongoing advancements in technology for recording brain and muscle signals, such as smaller setups with fewer channels, and wearable systems that deliver controlled peripheral stimulation, could further reduce the C-EPOCS system footprint. Addressing issues like trigger onset jitter at lower sampling rates may help overcome challenges associated with low-resolution signals. Further studies are needed to evaluate the feasibility of using a portable C-EPOCS system to measure afferent responses or deliver rehabilitation training outside the lab, such as in rehabilitation centers.

## Conclusion

This study outlines the customizations and evaluations to adapt the Cortical-EPOCS system into a portable platform for real-time brain-computer interface applications. The results demonstrate that the current component-heavy C-EPOCS setup can be effectively streamlined by using a single recording unit for both EEG and EMG acquisition– with minimal compromise to C-EPOCS applications– despite the reduced resolution of EMG signals. This setup can help maintain effective afferent excitation, supporting enhanced response stability and longitudinal tracking– both essential for cortical evoked response-based neurofeedback training.

### Ethical Considerations

The study was approved by the local Institutional Review Board at Stratton VA Medical Center, Albany, NY (Protocol #1584762). All data were collected with written informed consent. Participation was voluntary with ability to opt out at any time. The research was conducted in accordance with the principles embodied in the Declaration of Helsinki and in accordance with local IRB requirements.

## Conflicts of Interest or Disclosures

Authors do not have any conflicts of interest to disclose.

### Use of AI tools

Chatgpt model 4o was used for proofreading, to check grammar, tense, and punctuation, using an instruction ‘check English’. The updated text and the specific improvements were duly checked if they were included in the manuscript.

## Data Availability

The data can be made available per reasonable request and with required institutional data sharing agreement. Source data shown in the manuscript will be made available at OSF (doi:).

## Acknowledgements

We thank all the participants for their time and participation. We thank Dr. Wolpaw for his support for IRB and NCAN laboratory resources. We acknowledge the support in funding and resources from The National Institutes of Health (NIH-NIBIB) award P41 EB018783 (Wolpaw); NYS Spinal Cord Injury Research Board C37714GG (Gupta) and C38338GG (Wolpaw) and the Stratton Veterans Affairs Medical Center.

